# Rules Governing Perception of Multiple Phosphenes by Human Observers

**DOI:** 10.1101/302547

**Authors:** William Bosking, Brett Foster, Ping Sun, Mike Beauchamp, Daniel Yoshor

**Affiliations:** Department of Neurosurgery, Baylor College of Medicine, Houston TX; 2Department of Neuroscience, Baylor College of Medicine, Houston TX

## Abstract

Stimulation of single sites in primary visual cortex results in the perception of a small flash of light known as a phosphene. Little is known about how phosphenes from multiple electrodes can be combined to form perception of coherent patterns. Here we examine the percepts reported by human observers as various spatial configurations of 2 to 5 electrodes in visual cortex were stimulated simultaneously. When two electrodes were stimulated simultaneously, subjects reliably perceived either one or two phosphenes depending on the physical distance separating the electrodes. In cases where two phosphenes were perceived, they were located in the same visual field location as when the two electrodes were stimulated separately. Adding a third electrode produced similar results. In several subjects, we obtained combination of 4 to 5 electrodes that generated individual phosphenes when stimulated concurrently. Subjects were able to reliably discriminate between different multiple electrode stimulation patterns that were presented in random order. These results demonstrate that simple pattern information can be conveyed to subjects with surface electrodes spaced at millimeters apart on the cortex.

## Introduction

Electrical stimulation of the primary visual cortex (V1) evokes a simple visual percept know as a phosphene (Penfield 1947, Penfield & Rasmussen 1950). Soon after the discovery of this phenomenon, it was realized that this mechanism could form the basis for a visual cortical prosthetic (VCP)(Bosking et al. 2017a, Fernandes et al. 2012, Lewis et al. 2015, Lewis et al. 2016, Lewis & Rosenfeld 2016, Maynard 2001, Schiller & Tehovnik 2008, Tehovnik et al. 2009). The first generation of VCPs, which were implanted in blind subjects several decades ago, were partially successful (Brindley & Lewin 1968b, Dobelle & Mladejovsky 1974). Stimulation of single surface electrodes reliably produced phosphenes that were relatively small, and which were located in particular part of the visual field that corresponded to the location of the electrode in the map of visual space. Electrodes separated by approximately 2 to 3 mm on the cortical surface often generated distinct phosphenes (Brindley & Lewin 1968a). There were reports that simple patterns could be generated by stimulation of multiple electrodes (Dobelle et al. 1974), and that subjects could learn to discriminate different patterns (Dobelle et al. 1976), or even perform much more complex behaviors. However, these behavioral observations were poorly documented and quantified.

Only one series of experiments has used penetrating electrodes to evaluate the characteristics of phosphenes generated by electrical stimulation (Bak et al. 1990, Schmidt et al. 1996). These experiments demonstrated that less current is required to produce phosphenes with these electrodes, and that a smaller separation between electrodes was required for separate phosphenes. Unfortunately, however, these investigators were not able to examine pattern formation using multiple electrodes.

Recent work using electrical stimulation in epilepsy patients has examined how the size of phosphenes is related to the location of the electrode in the map of visual space (eccentricity) and to the parameters of the electrical stimulation train (Bosking et al. 2017b, Winawer & Parvizi 2016).

Currently, there is renewed interest in the development of VCP devices (Bosking et al. 2017a, Lewis et al. 2015, Lewis et al. 2016). Several groups are moving forward towards clinical trials either with subdural electrodes (Second Sight Medical Products 2015, Second Sight Medical Products 2016), or with arrays of penetrating electrodes placed on tiles that would float on the cortical surface (Lowery et al. 2015, Monash University 2017, Troyk et al. 2003, Troyk et al. 2005). However, we still know relatively little about how effective these electrode arrays will be in communicating pattern information to the subjects, and how to choose the most effect electrical stimulation protocols that should be used with these devices.

To begin to address this gap in knowledge, we have conducted extensive controlled testing with various spatial configurations of groups of 2-6 electrodes in early visual cortex in epilepsy patients. We found that concurrent stimulation of two sites separated by greater than (6-10 mm), or approximately one RF width, reliably generated two phosphenes. Configurations of three electrodes appeared to largely follow the same rules uncovered with stimulation of pairs, and we found that the average pairwise separation between electrodes within a triplet is correlated with the number of phosphenes that subjects perceive when three electrodes are stimulated at once. In addition, we found that we could stimulate up to 4-6 electrodes at once, and we documented subjects perception of up to 5 phosphenes. Our subjects could not only perceive patterns of phosphenes, but could reliably perform discrimination tasks using multiple patterns. However, one significant remaining issue is that our patients report that with concurrent stimulation of multiple electrodes they perceive a group of distinct spots, not a coherent object. These results highlight both the promise, and the complexity, of using electrical stimulation in VCPs.

## Materials and Methods

### Subjects

Electrical stimulation was conducted in patients (n = 8) with medically intractable epilepsy who had subdural electrodes implanted over various regions of the cortex for clinical purposes. Informed consent was obtained from all subjects, and the Baylor College of Medicine Institutional Review Board approved all procedures. The patients were typically kept in the epilepsy-monitoring unit for 4 to 14 days after the electrodes were implanted. Clinical monitoring continued uninterrupted during experimental sessions, which typically took place on first through fourth days post implantation.

### Electrodes

Although a variety of electrode types were implanted for clinical purposes, the only electrodes used for electrical stimulation in this study were research electrodes (platinum, 0.5 mm diameter) embedded in custom silastic strips (PMT Corporation, Chanhassen, MN). These custom strips were fabricated with the research electrodes positioned in the normally empty space between the larger standard clinical recording electrodes (platinum, 2.2 mm diameter, 1 cm spacing). The electrodes used in this study were located on the surface of the occipital pole, near the calcarine fissure (Figure 1A). This area is known to correspond to the primary visual cortex (V1), and other early visual cortical areas (V2, V3). Up to 16 electrodes were tested in each hemisphere. Three different hybrid clinical/research electrode strips were used with a variable number of research electrodes (8,12, or 16), but in all cases only research electrodes of the same type and diameter were used for electrical stimulation.

**Figure 1:**
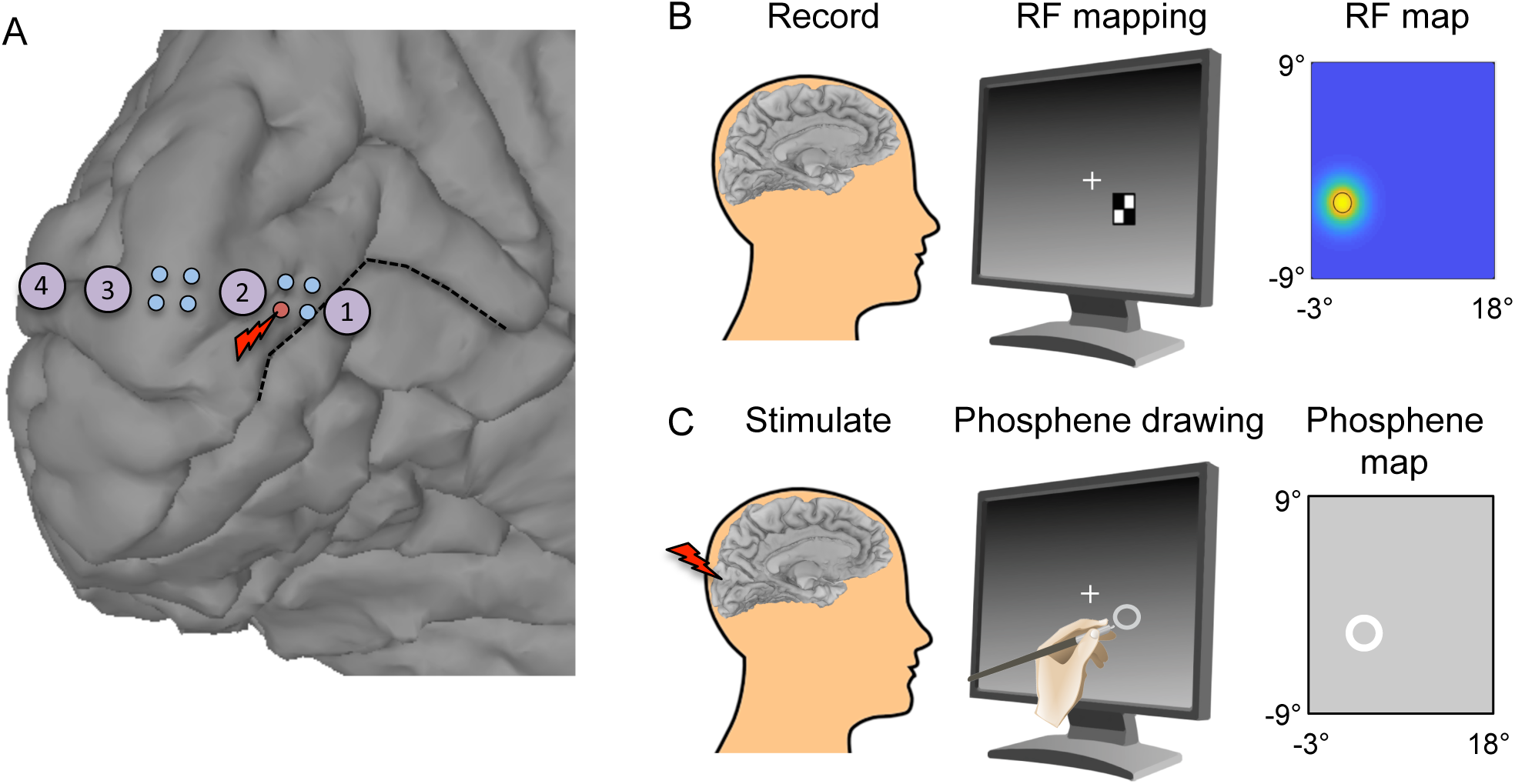
Recording and stimulation of subdural electrodes in human subjects. A) Location of a hybrid (clinical/research) electrode strip on the the medial wall of the occipital cortex in one subject. Research mini-electrodes (small blue circles, 0.5 mm in diameter, spaced at 2 mm intervals) were located in between standard clinical recording electrodes (red numbered circles, 3 mm diameter, spaced at 1 cm intervals). This electrode strip crosses the calcarine fissure (black dashed line) which identifies the location of primary visual cortex and the representation of the horizontal meridian. B) Receptive field mapping using checkerboard stimuli. Subjects performed a letter detection task at a central fixation point while checkerboard stimuli were presented in an array of locations on the screen. A 2D Gaussian curve was then fit to the responses to quantify the size and location of the RF. C) Phosphene mapping. Electrical stimulation was provided simultaneously to between 1-6 mini electrodes over early visual cortex while the subject fixated a small cross on touchscreen. Following the electrical stimulation the subject was asked to outline the phosphenes that they perceived using a stylus.

### Electrical Stimulation General

During all experiments, the patients remained seated comfortably in their hospital bed. A ground pad was adhered to the patient’s thigh, and all electrical stimulation was monopolar. Electrical stimulation currents were generated using a 16-channel system (AlphaLab SnR, Alpha Omega, Alpharetta, GA) controlled by custom code written in MATLAB (Version 2013b, The MathWorks Inc., Natick, MA).

We first screened all electrodes to determine which ones produced a phosphene when electrical stimulation was delivered. For this study, a phosphene is defined as a localized, brief, visual percept (commonly described as a flash of light). For each electrode, we typically began with a low current (0.3-1.0 mA) and gradually increased current on each trial until the patient reported a phosphene. If no phosphene was obtained with a current of 4 mA, then the site was considered unresponsive. A maximum of 4 mA was used, and the number of stimulation trials was limited, so as to maximize efficient use of time, and to limit the possibility of evoking seizures in our subjects. After screening, we conducted more detailed studies using different stimulation currents, and a variety of multi-electrode combinations.

During each stimulation trial, an auditory warning tone cued the patients to fix their gaze on a small cross on the touchscreen. This was followed by a second tone that indicated the beginning of the actual electrical stimulation period. Electrical stimulation consisting of a train of biphasic pulses (-/+), with 0.1 ms pulse duration per phase, was then delivered at a frequency of 200 Hz, with an overall stimulus train duration of 200 or 300 ms. Currents tested ranged from 0.3 - 4.0 mA resulting in a total charge delivered of 1.2 - 24 μC per trial.

### Phosphene mapping

Our phosphene mapping technique is illustrated in Figure 1C. Patients viewed an LCD touchscreen that was typically located 57 cm in front of them. The patient fixated on a cross on the display and electrical stimulation was administered following an auditory warning tone. The patients indicated whether they saw anything by verbal report, and then drew the outline of the visual percept (phosphene) using a stylus on the touchscreen. Multiple trials were typically conducted to allow the patients to precisely adjust the size and location of the outline. The subject was instructed to draw the shape as accurately as possible following the first stimulation trial, and then on subsequent trials they adjusted the scaling and location of the contour using a custom designed graphical user interface until it corresponded well to the phosphene they perceived. We instructed subjects in the use of the graphical user interface, but decisions about how to graphically depict phosphenes were made entirely by the subjects without interference. Electrodes that generated two completely distinct phosphenes (possibly representing an electrode spanning a sulcus) were encountered rarely, and were excluded from further testing and analysis. In some early cases, phosphene outlines were obtained by pencil drawings on paper on repeated trials rather than using the touchscreen. In these cases, a small cross was drawn on the paper for the subject to fixate.

### Analysis of phosphene maps

All phosphene drawings on the touchscreen were ellipses, and all phosphene drawings made on paper were fit with an ellipse. The center of the best-fit ellipse was taken as the center of the phosphene. We used (major diameter + minor diameter) / 2 as the measure of phosphene size. Phosphene size in degrees of visual space was calculated by using the standard formula for calculating visual angles: V = 2 arctan(S/2D) where V = visual angle in degrees, S = size of the object or stimulus in question, and D = the viewing distance. In our case: PSdeg = 2 ^*^ arctan(PScm/2D) where PDdeg = size of the phosphene in degrees of visual space, PScm = size of the phosphene drawn on the screen in cm, and D = distance from the eye to the screen in cm. For our typical screen distance of 57 cm, this resulted in 1 cm on the screen being equal to 1° of visual angle. The distance of the phosphene center from the fixation point (cm) was used to determine the eccentricity of the stimulation site (°) by multiplying by the same conversion factor (1°/cm).

### Electrode Localization

Electrode placements were guided solely by clinical criteria. Before electrode implantation surgery, two T1-weighted structural magnetic resonance (MR) scans were obtained. The two scans were aligned and averaged to provide maximum gray–white contrast using Analysis of Functional NeuroImages software (AFNI)(Cox 1996). Cortical surface models were constructed using the program FreeSurfer (Dale et al. 1999, Fischl et al. 1999), and visualized using the SUMA component of AFNI (Argall et al. 2006). After electrode implantation surgery, subjects underwent whole-head computed tomography (CT). The CT scan was aligned to the pre-surgical MR scan using AFNI, and all electrode positions were manually marked on the structural MR images. Subsequently, the electrode positions were assigned to the nearest node on the cortical surface model using AFNI.

## Model

Our model for prediction of phosphene size consists of two parts. The first part predicts the spread of cortical activity based on electrical current using a sigmoidal function, and the second part predicts cortical magnification factor using a previously published equation (Dougherty et al. 2003, Horton & Hoyt 1991). Finally, the diameter of activated cortex is multiplied by the inverse magnification factor to predict phosphene size.

### V1-V2-V3 Flat Map Model

We used a previously published set of transforms to model the V1-V2-V3 complex (Schira et al. 2009), and all cortical activation modeling and distance calculations were made after projecting electrode locations onto this map. To project the electrode location onto the map we used RF information if possible. If no RF information was available for that electrode then we used the center of the individual phosphene map. The predicted cortical activation diameter was obtained using the eccentricity of the electrode within the map, and using the magnitude of the current used for stimulation according to equations presented in our recently published manuscript (Bosking et al. 2017b). The scaling of the V1-V2-V3 complex was adjusted for each subject by adjusting the spacing between known landmarks, such as two of the clinical recording electrodes, to be equal to the nominal distance known to exist for those landmarks.

### Calculation of Distance on the Cortical Surface

We used a hybrid method to obtain the final best estimate of distance separating each pair of electrodes tested. If two electrodes were located close together on a cortical strip, and there was no evidence of a sulcus or fissure running in between those two electrodes, then the nominal distance on the electrode strip was used as the estimate of distance between the two electrodes. When it was clear that a sulcus did run in between two electrodes we instead used a distance measure derived from a cortical surface model that was made from MRI images for that subject. The cortical surface model was not used for the short distance calculations because we feared it would inject errors due to inaccuracies in the model generation and closest node placement on the surface model. These errors could be millimeters in magnitude, which is significant for our study when short electrode separation distances are involved, but much less so for long separation distances between electrodes of several centimeters.

### Calculation of Shift Indices

To measure how far apart two electrodes were in cortical space, or how far apart two RF or phosphenes were visual space coordinates, we used a metric we call a shift index. In visual space, the receptive field shift index is calculated by taking the distance between the two RFs in degrees and dividing by the average diameter of the two RFs in question. This results in a low shift index (<1) when RFs are highly overlapping, a moderate shift index (~=1) when RFs just barely overlap, and a large index (>1) for RFs that are separated by several times the diameter of the RFs. The analogous calculation was made for phosphenes (also in visual space), and for the predicted cortical activation (in cortical space).

#### Pattern Discrimination Experiments

The discrimination experiments were 2AFC or 3AFC with a single stimulus interval. Trials with different patterns were presented in random order. On each trial, a tone was played followed by the electrical stimulation. The subject was instructed to fixate on a small cross on the screen during the brief presentation of the electrical stimulus. After the electrical stimulation the subject gave a verbal report of which of the patterns they perceived on that trial. In some cases, a sheet of paper with the possible pattern choices was placed in front of the subject below the visual stimulus screen to remind the subject what the possible choices were.

## Results

We conducted electrical stimulation in epilepsy patients with normal vision (n =8) while they were hospitalized with cortical surface grids and strips for clinical monitoring purposes (Yoshor et al. 2007). Electrodes were located near the occipital pole and near the calcarine fissure on the interhemispheric surface (Figure 1A). We used research mini-electrodes (0.5 mm in diameter) for both recording and electrical stimulation. Three different electrode configurations were used with 8, 12, or 16 research electrodes located in the space between the normal clinical recording electrodes.

In each subject, we initially screened all of the electrodes in the early occipital cortex to determine which ones produced phosphenes when stimulated in isolation with low currents. Electrical stimulation trains were 200 – 300 ms in duration, 200 Hz, 0.1 ms pulse width, biphasic, monopolar. As previously reported (Bosking et al. 2017b, Murphey et al. 2009, Murphey et al. 2008), using these stimulation parameters, threshold currents for producing single phosphenes were typically quite low (mean = 0.8 in previous work). We then tested various combinations of 2-6 electrodes stimulated concurrently.

In 6 subjects, we tested concurrent stimulation of as many different pairs of electrodes as possible (224 total). To measure the cortical surface distance between two electrodes we used a hybrid approach. When electrodes were separated by a short distance on the electrode strip, and it appeared that the two electrodes in question did not lie on opposites sides of a cortical sulcus or fissure, we used the nominal distance apart between the two electrodes on the electrode strip. When the two members of a pair appeared to lie on opposites side of sulcus, we used a cortical surface model based on MRI measurements to estimate the actual distance along the cortical surface (see methods). As expected, subjects tended to perceive two phosphenes when the distance separating the two electrodes was large (>6-10 mm; Figure 3A). In some cases when only one phosphene was perceived it seemed clear that the subject perceived the stimulation of only one of the two electrodes, rather than a blending together of activity from the two electrodes. This was easy to demonstrate since the subject drew the single phosphene obtained with simultaneous stimulation at a location very close to the position of one of the two locations where phosphenes were perceived when the electrodes were stimulated independently.

We tested a number of other factors to see if they were effective in allowing us to predict the number of phosphenes that a subject would perceive when a particular pair of electrodes was stimulated. In addition to the cortical surface distance separating the two electrodes, the two other factors that were most effective were a receptive field shift index (Figure 3B), and a cortical activation index (Figure 3c). The receptive field shift index simply calculated the separation in visual space between the receptive fields of the two electrodes (°), and divided this value by the average receptive field width (°) for the two sites, resulting in a dimensionless index. This measure gives a value of less than one when the RFs for two sites are overlapping, and a value greater than one when the two receptive fields are shifted in visual space by more than one RF diameter. The cortical activation shift index was calculated using a similar procedure. For each electrode in the pair, we predicted the center and diameter of the expected activity pattern within the map of visual space (Figure 2; Figure 6). The center of the activation pattern was predicted using the receptive field center for that electrode if that information was available, and was based on the center of phosphenes obtained when electrodes were stimulated independently when no RF information was available. The width of the predicted activation pattern was calculated based on the eccentricity of the electrode within the map of visual space, and on the magnitude of the current injected (Bosking et al. 2017b). We calculated the separation between the centers of the two predicted activity patterns within the cortical map of visual space (mm) and divided by the average width of the activation patterns (mm). This results in a cortical activation shift index that is above one if the two activity patterns do not overlap in the cortex, and less than one for activity patterns that are expected to overlap significantly. The idea behind this measure is that when there are two clearly separate peaks of activity in the cortex we would expect the subject to see two phosphenes, and when the two activity patterns merge into one continuous region of activity we would predict that the subject should be more likely to see only one phosphene, that is perhaps larger in size. The predicted cortical shift index appeared to be a sensitive predictor of the number of phosphenes observed with pair stimulation (Figure 3C).

**Figure 2:**
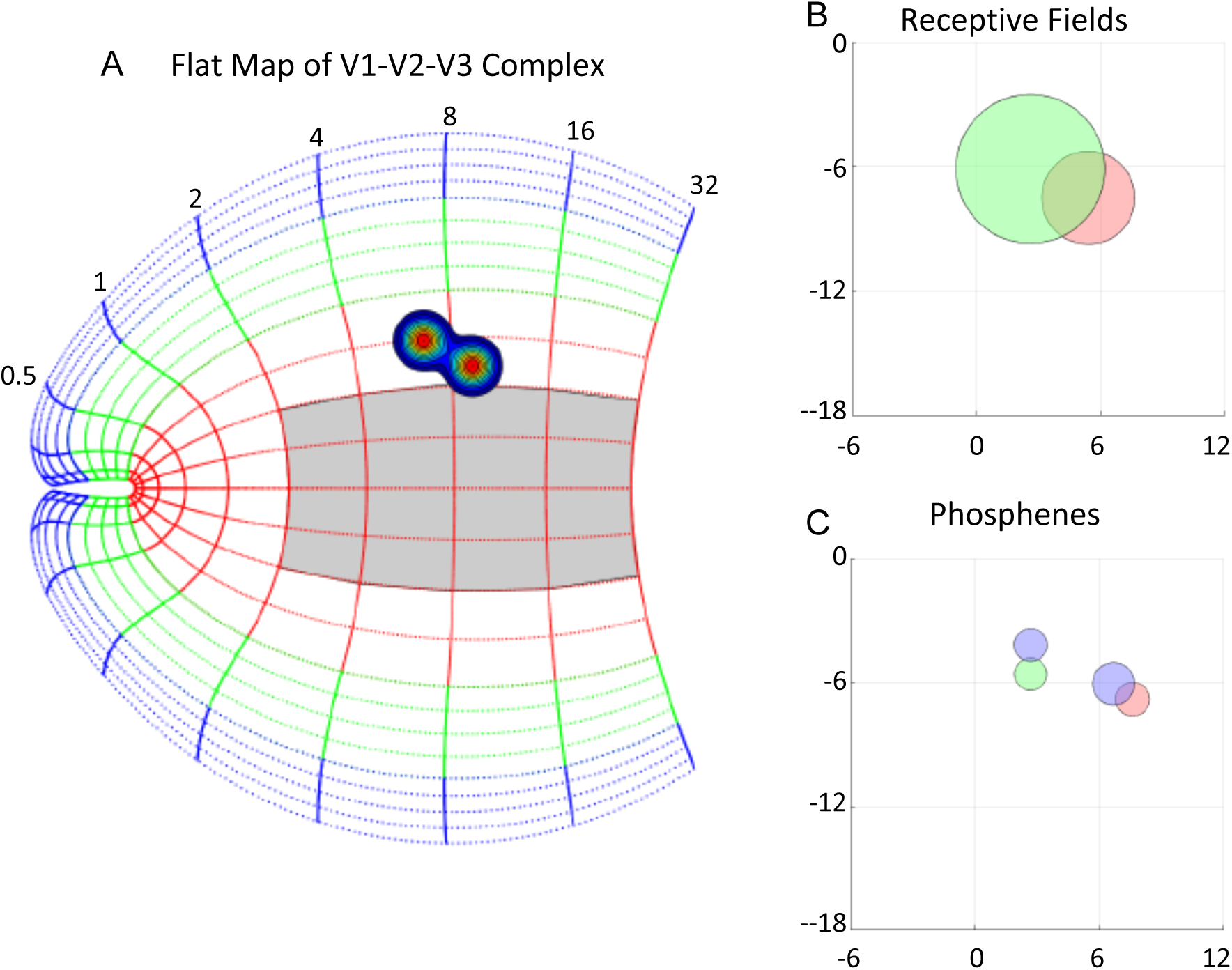
Example data for stimulation of one pair of electrodes in one subject. A) The estimated location of the stimulated parts of early visual cortex are indicated on a flat map of the V1-V2-V3 complex. Location of the center of each activation area is based on the RF centers for the corresponding electrodes when available, or using the location of phosphenes that were obtained with single electrode stimulation when RFs were not available. The size of the activated cortex was predicted according to the eccentricity of the electrode in the map of visual space, and the current that was used for stimulation (Bosking et al, 2017). B) Receptive field location and size for the same two electrodes used for stimulation. C) The location of phosphenes obtained with single electrode stimulation (red and green circles) and when the pair of electrodes was stimulated simultaneously (blue circles).

**Figure 3:**
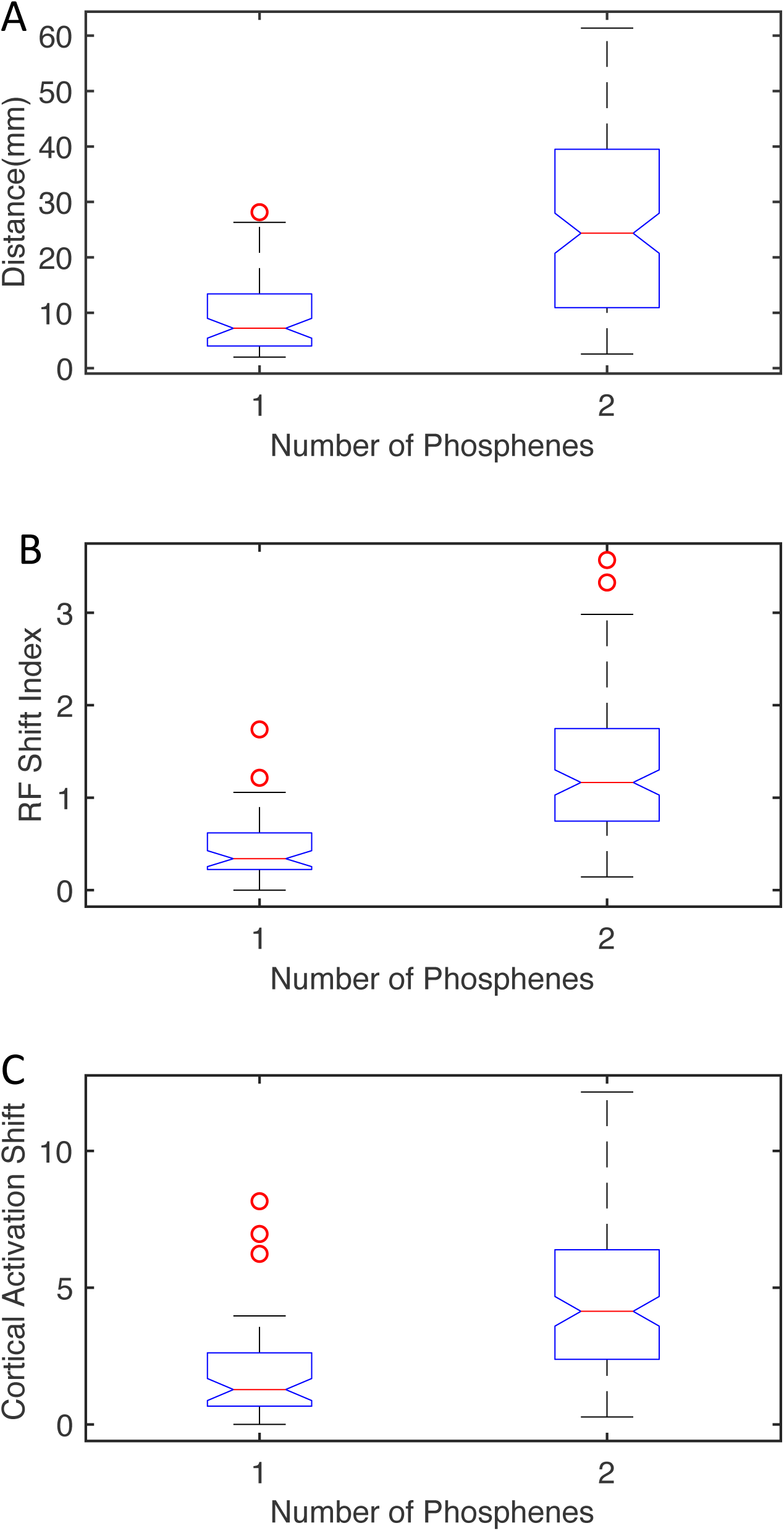
Summary data for stimulation of pairs of electrodes. A) Separation distances between the pair of two electrodes at which one (left) or two (right) phosphenes were obtained when the electrodes were stimulated simultaneously. On each boxplot, the red line indicates the median, the bottom and top edges of the box indicate the 25^th^ and 75^th^ percentiles, and the whiskers extend to cover approximately 99% of the data. Outliers are indicated by red circles. B) Receptive field shifts between electrode pairs that generated one or two phosphenes when stimulated simultaneously. A receptive field shift value of 1 indicates that the RFs for the two electrodes were approximately non-overlapping. C) Shift in predicted cortical activation for electrode pairs that produced one or two phosphenes when stimulated simultaneously. A value of 1 indicates that we would predict that the two activation pattern should not overlap, or will overlap by only a small amount.

There was a trend for phosphenes associated with a particular electrode to be slightly smaller in size when the electrode was used as part of a pair (Figure 4). In general, phosphenes perceived during concurrent pair stimulation were in near the same location in visual space as those obtained with individual stimulation of each electrode (Figure 2; Figure 6). In at least one subject, however, there was a strong trend towards greater separation between the two phosphenes generated by the pair of electrodes when the pair was stimulated simultaneously as opposed to independently (Figure 5).

**Figure 4:**
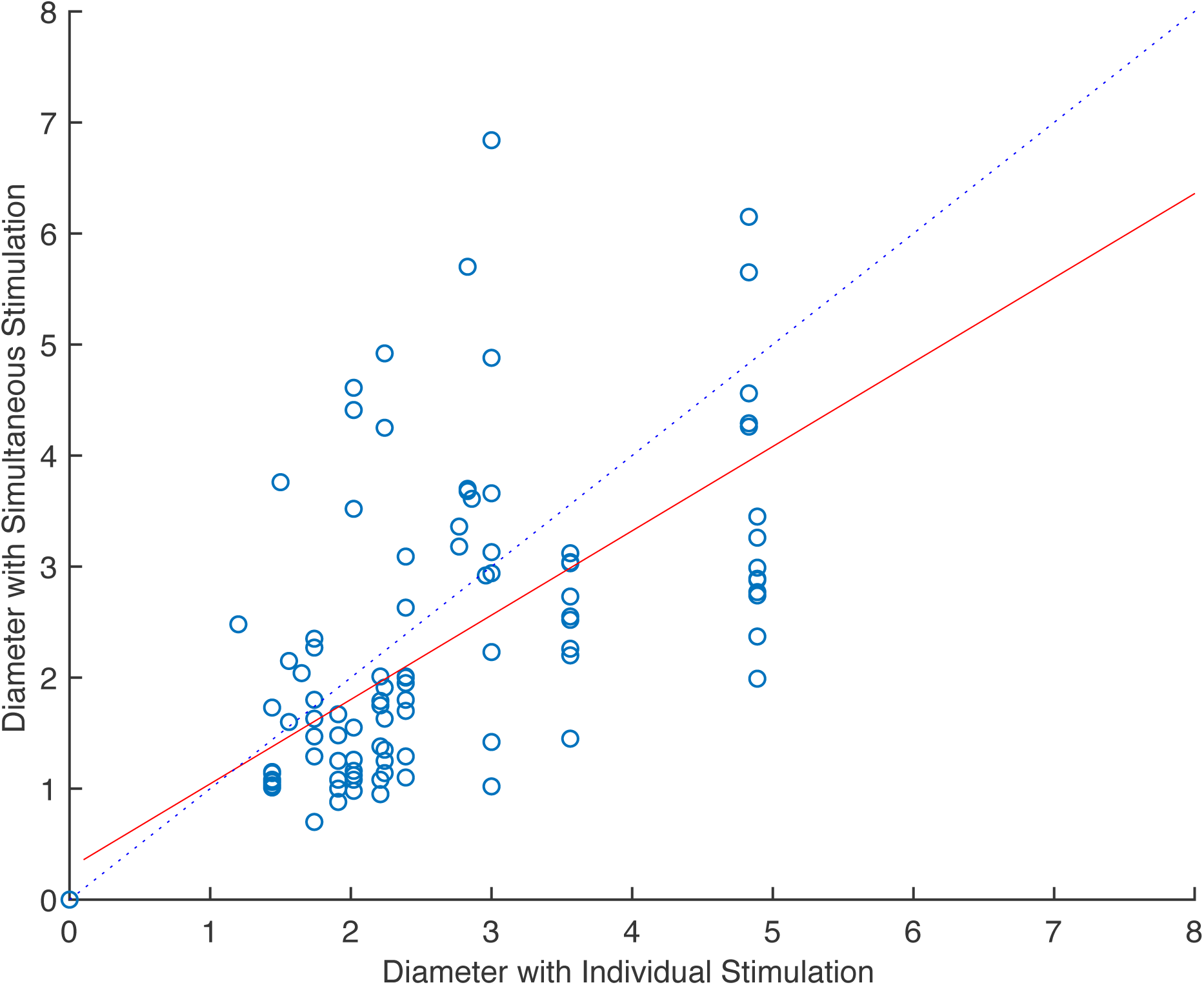
Diameter of phosphenes when pairs of electrodes were stimulated independently or simultaneously. Each data point corresponds to one pair of electrodes. The blue dashed line indicates the point at which equal sized phosphenes would be obtained in the two stimulation conditions, and the solid red line indicates the linear fit to the data.

**Figure 5:**
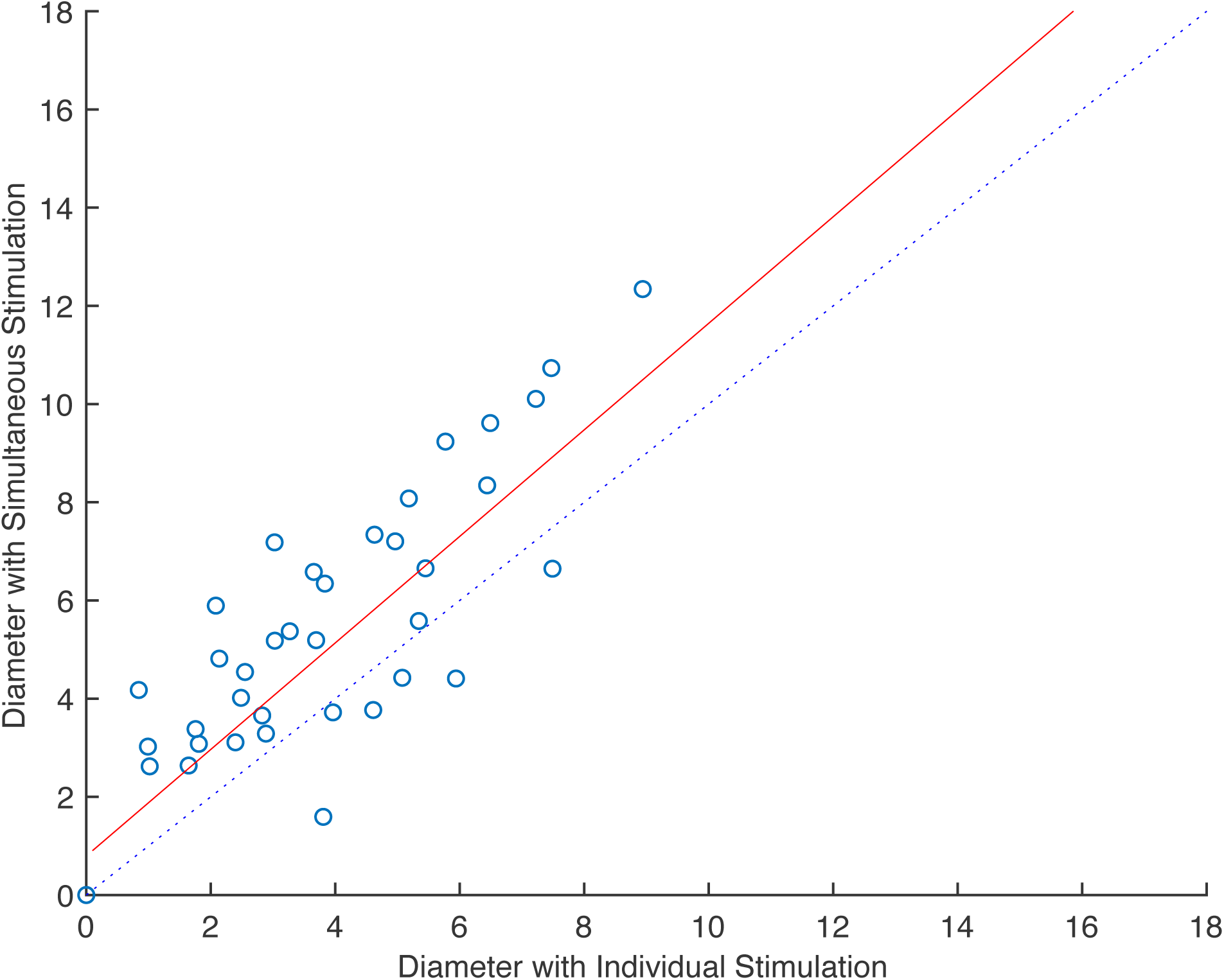
Separation between plotted phosphenes when electrode pairs were stimulated simultaneously or independently. Each data point corresponds to one pair of electrodes. The blue dashed line indicates the point at which equal separation between the two phosphenes would be obtained in the two stimulation conditions, and the solid red line indicates the linear fit to the data.

Similar results were obtained when we tested concurrent stimulation of three electrodes (n = 4 subjects; 63 triplets; Figure 6). To see if distance separating the electrodes in the triplet could still be used to predict how many phosphenes the subject observed, we calculated the pairwise separation distances between each of the three possible pairs within the group of three electrodes (1-2, 1-3, 2-3). We found that this measure was strongly correlated with the number of phosphenes perceived by the subject. One phosphene was perceived when the electrodes had a relatively small average pairwise separation (11.28 mm), two phosphenes were associated with a higher average pairwise distance (23.52 mm), and three phosphenes were perceived when the average pairwise distance was higher still (37.17 mm).

**Figure 6:**
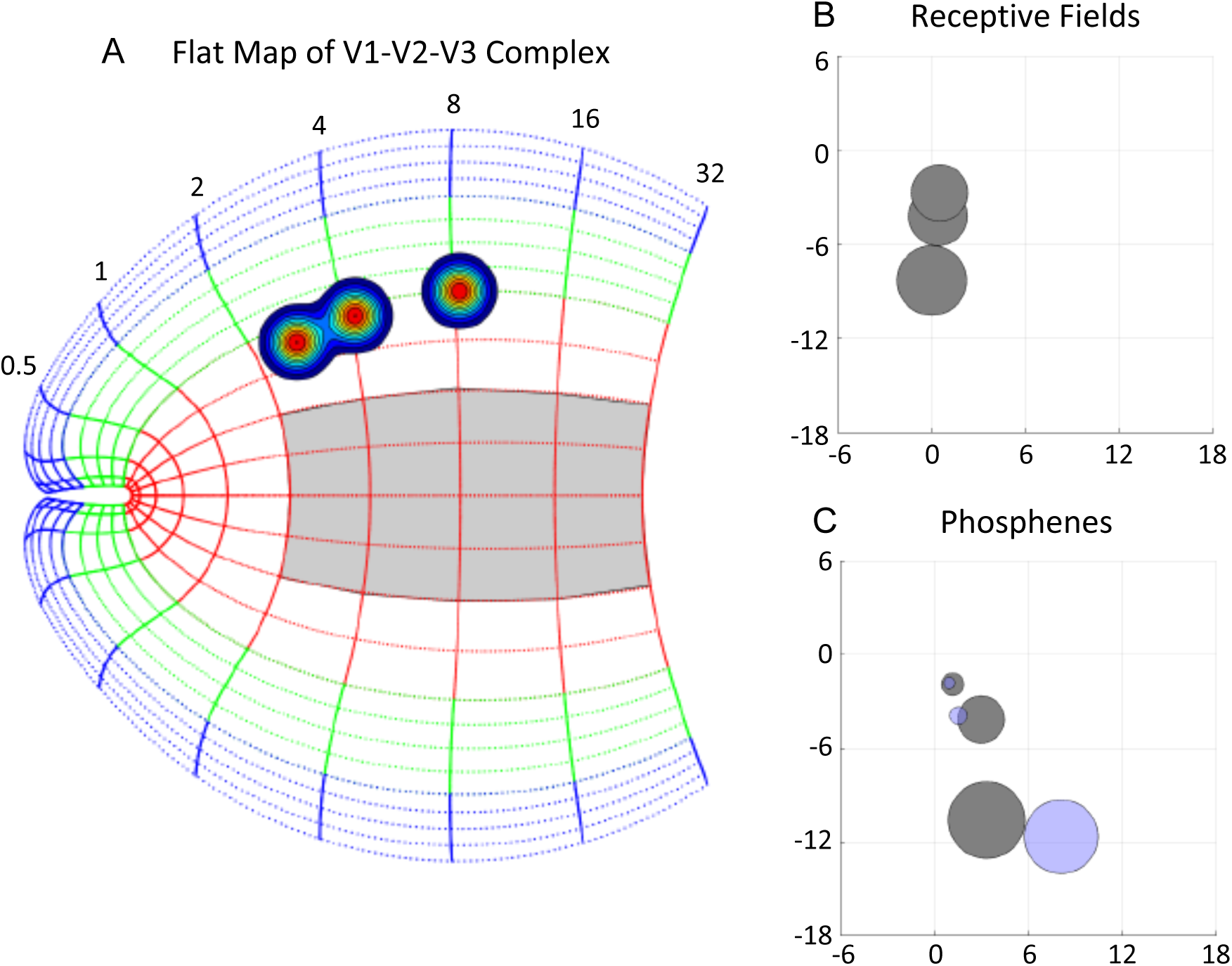
Example data for stimulation of one set of three electrodes in one subject. A) The estimated location and size of the activated regions of early visual predicted using the same methods as described for Figure 2. B) Receptive field location and size for the same three electrodes used for stimulation. C) The location of phosphenes obtained with single electrode stimulation (gray circles) and when the pair of electrodes was stimulated simultaneously (blue circles).

**Figure 7:**
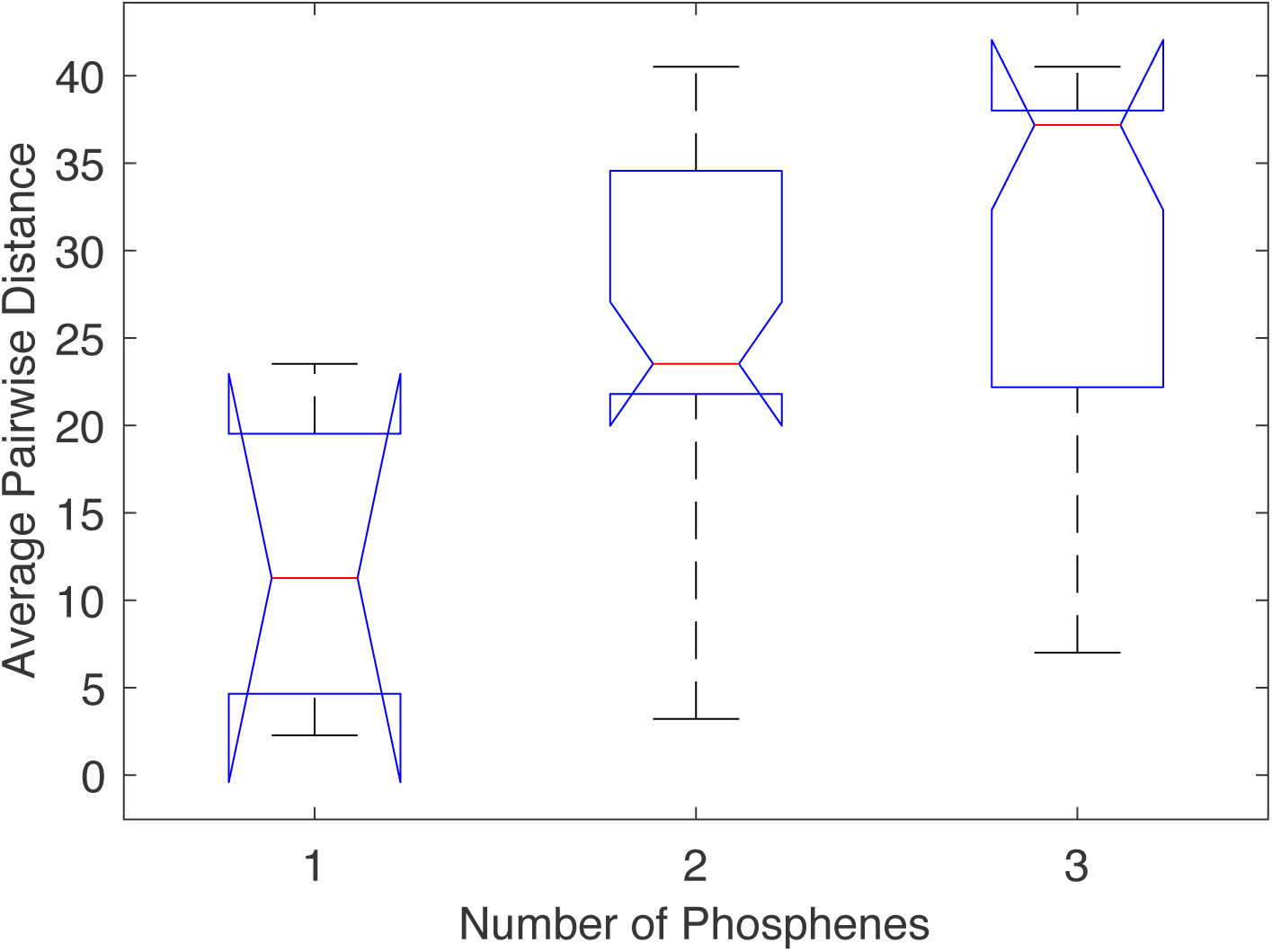
Summary data for stimulation of triplets. Average pairwise distance between the three electrodes in the set is shown for groups that produced one, two, or three phosphenes when stimulated simultaneously. Boxplots indicate the median and 25^th^ and 75^th^ percentiles as in Figure 3.

In two subjects we conducted two alternative or three alternative forced choice tasks (2AFC, 3AFC). On each trial, the subject was presented with one of the two (or three) multi-electrode patterns under examination. The subject gave a verbal report as to which of the possible patterns they perceived on that trial. In eight 2AFC experiments from the two subjects the average behavioral performance was 80.4%, and in two 3AFC experiments from one subject the average performance was 50.9%

## Discussion

### Overview

We performed an extensive set of testing of multi-electrode stimulation of early visual cortex in humans. Our results are on the one hand encouraging, since they clearly demonstrate that stimulation of specific patterns across the retinotopically mapped areas in early visual cortex can result in the perception of phosphenes in the expected locations, and that subjects can use these patterns to perform discrimination. On the other hand, our results also reveal some of the current limitations of our ability to deliver patterned stimulation to visual cortex.

### Cortical Separation Required for Perception of Two Phosphenes

Our estimate from this study of the separation required between two electrodes for the subject to perceive two phosphenes (6-10 mm, Figure 3A), is somewhat higher than we would have predicted from our recent study (Bosking et al. 2017b), our from earlier work (Brindley & Lewin 1968a). We reported that phosphene size could be predicted accurately if the factors of eccentricity within the visual field and magnitude of current injected were modeled. Specifically, we found that phosphene size saturated with increase in current above a moderate level of 1-2 mA, and that this appeared to be consistent with activation of a region of cortex approximately 5-6 mm in diameter. So, in this study we might have predicted that cortical separation distances of 5-6 mm would be sufficient for the subject to see two phosphenes with concurrent pair stimulation. Although this did happen sometimes, often the subject only reported one phosphene. Our earlier study may have underestimated the full range of cortical interactions that are evoked when electrical stimulation is delivered. For example, weak sub-threshold activation from the two electrodes could combine to bring further regions above threshold and result in a larger overall area activated than would have been predicted by the single electrode experiments.

### Prediction of Phosphene Combination by RFs

We found that using a RF shift index provided one way to predict whether the subject would perceive one or two phosphenes with pair stimulation. Before this round of experiments we might have predicted that the individual phosphene drawings made by the subject would provide the best estimate of the outcome of pair stimulation. Operationally, however, the RF shift index appears to provide better prediction capability, at least when a low number of trials is used for the phosphene mapping. The RF estimates are likely to be substantially more accurate than predictions based on phosphene maps due to both the number of trials used to obtain the RFs, and potential eye movements or offsets during phosphene mapping. In the best cases, separation on the cortical surface, RF shift index, and shifts in phosphene maps between the two electrodes are, of course, all highly correlated.

### Other Interactions

We noted that in at least some cases there was a strong tendency for phosphenes to be located further apart during simultaneous stimulation than in the independent stimulation condition, and that phosphenes often appear to be smaller in the concurrent condition. These effects may be additional clues that our single electrode experiments do not predict the full spatial radius, or the full complexity, of the interactions that take place when multiple electrodes are stimulated concurrently. One interesting possibility is that inhibitory or suppressive effects might have a larger interaction radius than excitatory effects. Experiments combining stimulation and fMRI in monkeys have demonstrated that large-scale inhibition can be evoked by electrical stimulation of the cortex (Logothetis et al. 2010), and that this can significantly effect signal propagation in the visual cortex.

### Implications for Visual Cortical Prosthetics

Our results confirm the precision with which we can input information into the early visual cortex because of the precise retinotopic maps in these areas. We show that subjects can use simple pattern to perform discriminations with little to no training in the less than optimum hospital environment. However, it is clear that to make spatial forms that appear more coherent to the subject, and which resemble shapes as complex as graphemes, that we probably need to rethink our stimulation strategy. We need to develop stimulation strategies that do not simply view electrodes on the cortical surface (or those that penetrate into the brain) as the equivalent of pixels on a monitor. We are currently exploring ways the stimulation patterns can be dynamically modulated so as to convey more coherent and precise spatial information in our epilepsy subjects. And of course it will be necessary to examine cortical interactions in blind patients as well since the nature of these interactions in the visual cortex may have changed as a result of blindness.

## Acknowledgements

Funding: NIH EY023336.

BLF is supported by an NIMH Career Development Award R00MH103479

## Notes

**Conflict of Interest** William Bosking receives funding from Second Sight corporation, the manufacturer of the Orion visual cortical prosthetic device.

